# Framework for assessing the risk to a field from fraudulent research

**DOI:** 10.1101/2024.12.23.630160

**Authors:** Chaoqun Ni, B. Ian Hutchins

## Abstract

Concerns over research integrity are rising, with increasing attention to potential threats from untrustworthy authors. We established a framework to gauge the potential negative influence of researchers potentially engaged in misconduct. The field of Alzheimer’s Disease (AD) research has been a focal point of these worries. This study aims to assess the risk posed by questionable studies or individuals potentially engaging in fraudulent science in research by examining citation relationships among papers, taking AD research as an illustrative example. Analysis of citation network structure can elucidate the potential propagation of misinformation arising at the author level. Our analysis revealed that there aren’t any single authors or papers whose citation connections jeopardize a major portion of the field’s literature. This indicates a low probability of single entities undermining the majority of works in this area. However, our findings suggest that attention to research integrity of the most influential scientists is warranted. Some scientists can reach a sizable minority of the literature through citations to their work. Emphasizing oversight of the integrity of these authors is crucial, given their influence on the field. Our study introduces an analytical framework adaptable across various fields and disciplines to evaluate potential risks from fraudulence.

## Introduction

Scientific publications rely on the practice of citation as a means to acknowledge and build upon previous research. Thus, citations in scientific literature are crucial for acknowledging intellectual contributions and tracing downstream impact. However, this process depends on accurate reporting of experimental results by scientists acting in good faith. Scientists engaging in fraudulent research practices, such as data fabrication and dredging, pose risks to the advancement of knowledge in their field. Fraudulent research may mislead researchers and waste resources, ultimately slowing down the progress of science and hindering the development of new knowledge. They may also erode public trust in science. This can undermine public confidence in science and make gaining support for scientific research and initiatives more difficult. Because biomedical research often informs life-or-death health decisions, fraudulence in this domain also poses a risk to public health (Marcus, 2018).

One field where research misconduct has received much attention recently is the Alzheimer’s Disease (AD) literature. A recent investigation in *Science* of a prominent neuroscientist working in AD research noted the potential scope of damage done by this alleged research misconduct (Weiland, 2022). This report, entitled “Blots on a field?”, called into question the entire amyloid hypothesis literature, noting that more than 2,300 articles had cited the work in question, which was published in 2006 (Weiland, 2022). Researchers, clinicians, and patients may have been relying on the reported findings to inform the development of new treatments or to improve patient care. If the allegation were proven true, it might have significant negative consequences. It could mean that those efforts were based on flawed or inaccurate information, and it could set back progress in the field. Other reports of repeated misconduct findings in neuroscience raise the importance of this issue (Kozlov, 2023).

Because scientific advances rely heavily on past discoveries, understanding the vulnerability of scientific knowledge networks to researchers potentially fabricating results is crucial for assessing the risks posed by research misconduct. Scientific knowledge networks are generally thought to conform to a “small-world” network structure, whereby large components of an information network can be influenced by a single hub like a well-cited paper (Bornmann et al., 2015). For this reason, it is theoretically plausible that a single fraudulent researcher could indeed cast doubt on a majority of a field of work, if the latter built upon and cited the author’s prior fraudulent work. Addressing this concern requires a high-level view of the scientific literature, as well as comprehensive and high-resolution data about authors, articles, and citations. Until recently, such data were not broadly available (Hutchins, 2021), but recent advances in data collection, processing (Hutchins et al., 2016), and open science (Hutchins, Baker, et al., 2019) have now made such questions tractable to answer at scale (Hutchins, Davis, et al., 2019). By analyzing the structure of the scientific literature and citation networks, it is now possible to assess the potential reach of individual researchers and their impact on the field, and therefore, the potential scope of damage if their work were fraudulent. This analysis provides insights into the extent to which a single researcher can cast doubt on a significant portion of a research field.

In this study, we develop a risk assessment framework for evaluating the potential reach of individual papers as well as scientists in a field of research through scientific knowledge networks. We characterize the AD literature from a high level. We investigate the potential impact that individual malicious papers would have on research in a field based on citation networks. We also analyze the network structure from an author-centric perspective, and determine the reach that individual authors have in the existing AD knowledge graph through subsequent citations. We find that most AD authors reach less than 1% of the literature through citations. Some scientists do reach over 5% of the literature, but these constitute far less than 1% of authors. Despite the small-world network architecture of citation graphs, we find that there are no author-level network hubs that can reach the majority of a field of research through first-order citations. However, attention to the research integrity of the most productive and influential scientists is warranted, since some can reach a sizable minority of the literature through direct citations to their work.

## Results

### Overview of the Alzheimer’s research enterprise

The US is estimated to have 6.7 million AD patients as of 2023 (Alzheimer’s Association, 2023), imposing massive economic and societal challenges. Accordingly, the AD research enterprise has increased substantially in the past decades (**Figure** 1). The number of publications on AD has increased steadily from a small number of papers per year in the 1980’s to several thousand per year by 2020 (**Figure 1.A**). The number of authors contributing to the AD literature has risen even more sharply (**Figure 1. A**), consistent with an increased number of authors per paper in biomedicine (Fortunato et al., 2018). These trends may accelerate in the future, as projects and funding from the National Institutes of Health (NIH) for Alzheimer’s research have increased exponentially in recent years (Figure 1. **B**). Publications lag funding, so the effect of this increased investment will not be visible until several years have passed.

**Figure 1.**
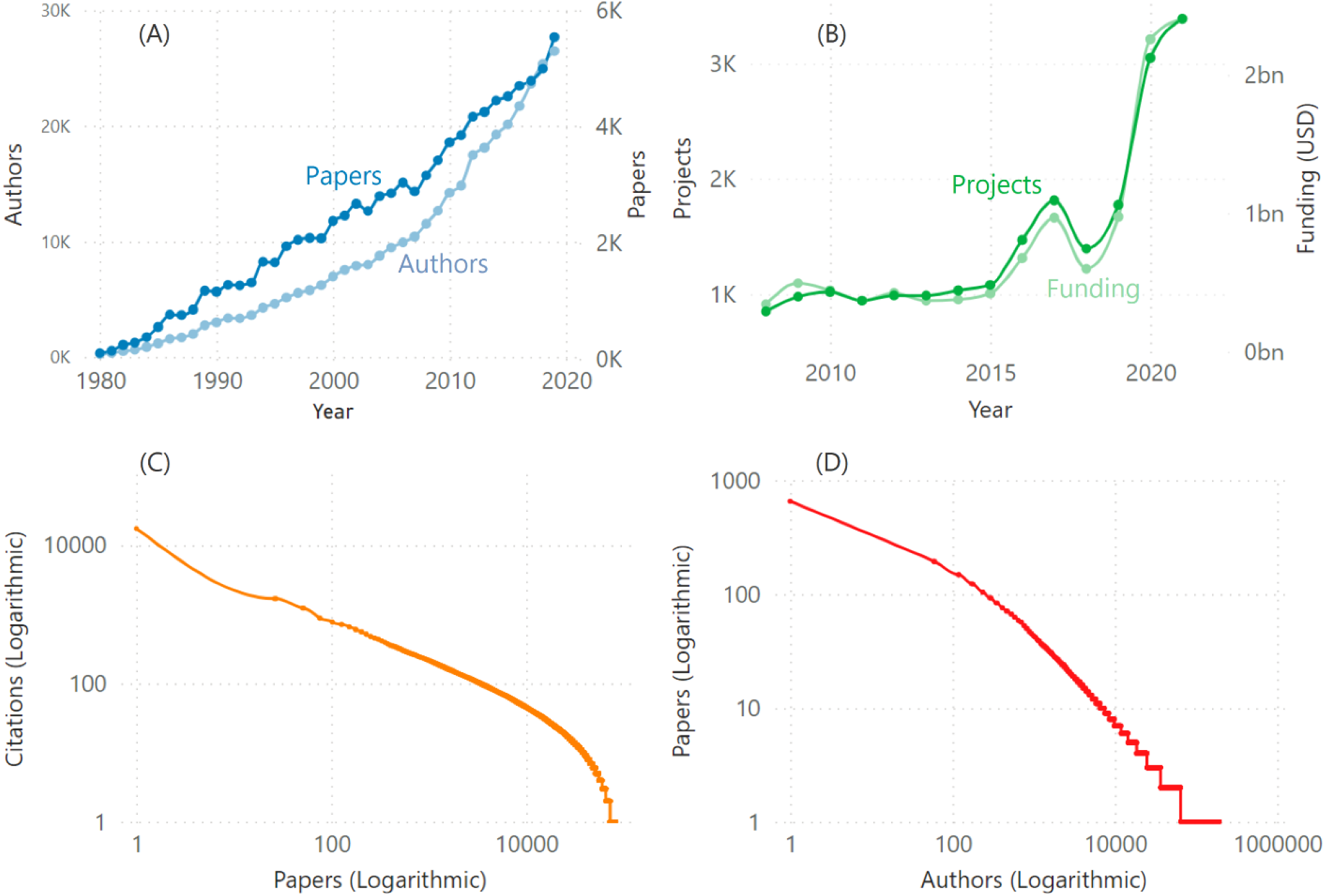
An overview of the AD research enterprise. (A) Number of publications and associated authors by time. (B) Number of NIH projects and amount of funding (in US dollars) by time. (C) Distribution of citations by paper; shown in logarithmic scale. (D) Distribution of the number of papers by author; shown in logarithmic scale

Given the increasing fraction of people afflicted by AD (Paroni et al., 2019) and the increased support from government funding agencies, the number of papers in this literature is likely to transition to a super-linear rate of advance in the coming years. In our analytical sample, the core Alzheimer’s literature consists of 112,720 publications contributed by 193,077 individual authors, creating 2,051,603 citation relationships. The publication quantity by individual authors follows an approximate power law distribution (Figure 1. C), implying that a limited number of authors dominates the production of knowledge in Alzheimer’s research. Citations to these papers also seem to center around a small proportion of papers in the sample (Figure 1. D): 10% of AD publications received 57% (n = 112,720) of AD citations. These findings indicate the potential impact of a small set of prolific authors and highly cited papers, which leads to further questions of how papers by those authors, if later found to be fraudulent, would contaminate the field if deemed unreliable, compared with other papers and authors.

### Risk analysis of individual papers

The AD literature has garnered a significant number of citations, reaching a total of 4,694,679. Within this corpus, 2,051,603 citations originated from the core AD research corpus (Methods). Notably, the distribution of citations to AD papers follows an approximate power law pattern. Specifically, a mere 10% of the papers accounted for a substantial 57% of the total citations. This observation suggests that citations tend to concentrate on a small group of publications, indicating a potential concentration of influence (Peng, 2015; Wang, 2014; Wang et al., 2008). This finding echoes previous research on the Mattew Effect of citation practices in many disciplines. Other less-cited papers, if subsequently called into question, might contaminate knowledge flow through the field via their citation network. Therefore, we calculated the shortest path in the citation network of AD papers to quantify the reach size of an individual AD paper in the knowledge network. The average shortest path in the network created by the citation relationship among AD papers is 4.4 (Figure 2. A), suggesting that knowledge from one AD paper will need to navigate through about 4.4 other papers in order to have citation influence on a target paper. Specifically, the 2,300 citations to the paper being investigated for potential research misconduct (Weiland, 2022), although significant in absolute terms, represent a small 2% (n = 112,720)fraction of all the literature.

**Figure 2.**
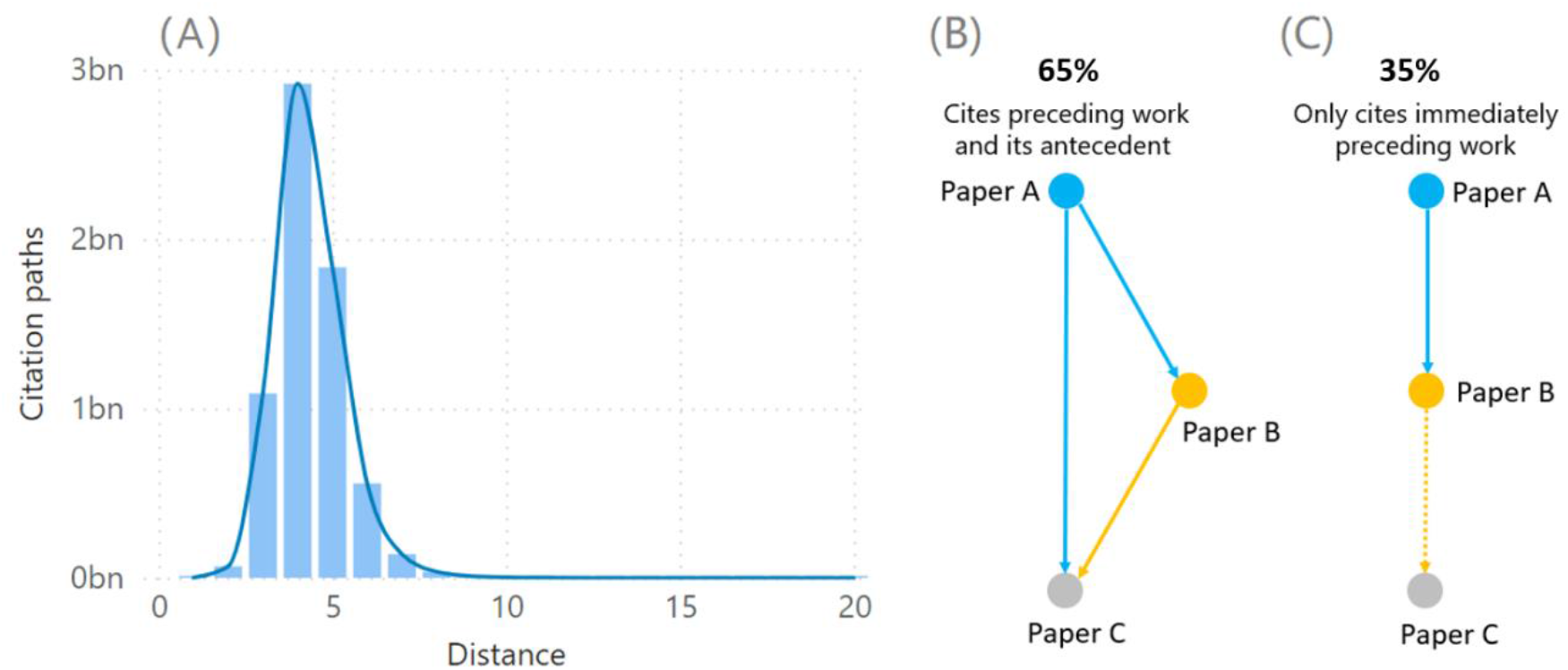
Network properties of AD paper citation relationships. (A) The distribution of the shortest path for papers in the AD paper citation network. (B) The tendency of authors to acknowledge both recent and antecedent work with citations, which forms a triangular local network structure (65% of citations). C: Linear citation chains when authors only cite recent work and ignore antecedent work (35%).

First-order fraudulent knowledge transmission when one paper builds upon another is not the only potential problem. Knowledge corruption might propagate to second-order citations (those citing a paper that cited a fraudulent paper). Biomedical research articles typically cite an average of 30 other papers (Hutchins et al., 2016). However, not all of these citations contribute to significant knowledge transfer that impacts the inception, design, or execution of the primary experiments (Hoppe et al., 2022; Hoppe et al., 2023; Teplitskiy et al., 2022). In other words, it is uncertain whether references-of-references, also known as “indirect citations,” more frequently convey substantial knowledge transfer or if they predominantly represent a relatively less significant form of rhetorical knowledge transfer.

Whether indirect citations can be considered substantive knowledge transfer depends on how researchers tend to cite previous papers. If researchers commonly cite both the most recent paper and the antecedent paper as the original source of knowledge, potential inaccuracies in knowledge transfer would likely be documented through both direct and indirect citations, forming a citation triangle (Figure 2.B). However, if researchers mainly cite the most recent article and neglect references to important antecedent work, we would predominantly observe linear citation chains.

To gain a better understanding of the impact of individual publications, we conducted an analysis of indirect citations among papers to gain a better understanding of the potential impact of individual publications. Specifically, the concept of indirect citation is based on the idea that when a paper (Paper A) cites the preceding work (Paper B), which in turn cites another work (Paper C) that Paper A did not directly cite, paper A provides an “indirect citation” to paper C. In some cases, Paper A may also directly cite Paper C. This means that if Paper C is deemed questionable, it, in principle, could corrupt Paper A in both scenarios, although this would be expected to be more likely in a citation culture where citation triangles are uncommon and predominantly linear citation chains are the norm. As such, we also assessed the likelihood of AD papers in our sample citing both the preceding and antecedent works versus only citing the immediately preceding work. We found that 35% of papers only cited immediately preceding work, while 65% cited their immediately preceding work and at least one antecedent simultaneously, significantly higher than a randomized network (p < 2.2e-16, n = 2,051,603, Fisher’s two-sided exact test). This result indicates a normative culture of citing both the more recent work in the AD literature as well as important preceding work. Therefore, indirect citations are less likely to account for the primary knowledge transfer in a field because antecedent work also appears as a direct citation in the majority of cases.

### The risk of citing individual authors

It is not uncommon for a series of works, rather than just a single piece, by the same author to come under scrutiny regarding their validity and reliability. There have been numerous examples (Kozlov, 2023; Normile, 2012; “Physicist found guilty of misconduct,” 2002) where doubts have been raised about the credibility of a collection of works by specific authors. The investigation findings published in Science (Weiland, 2022) raise an important question: how vulnerable is this field of AD research to potentially fraudulent information from individual scientists rather than papers? In order to gain insights into the potential impact that individual authors may have on the research community, we conducted an analysis of their author-level citation reach. We excluded from this analysis authors who only published a single first- or last-author paper in the AD literature or only middle-author papers since these authors may be ancillary to the field. We examined the extent to which other researchers cite their works to assess the potential risk associated with relying on the research output of these authors.

The papers among the 22,941 authors who published multiple first- or last-author AD research also shows a highly skewed distribution, similar to the article-level data (Figure 3), with about 10% of authors contributing to 51% of the papers in our dataset. Likewise, about 70% of citations (of n = 2,051,603) go to 10% of authors (n = 22,941). Therefore, the risk associated with potential fraudulent activities of a highly productive or highly cited author would be different from those of a less productive or less well-cited author.

**Figure 3.**
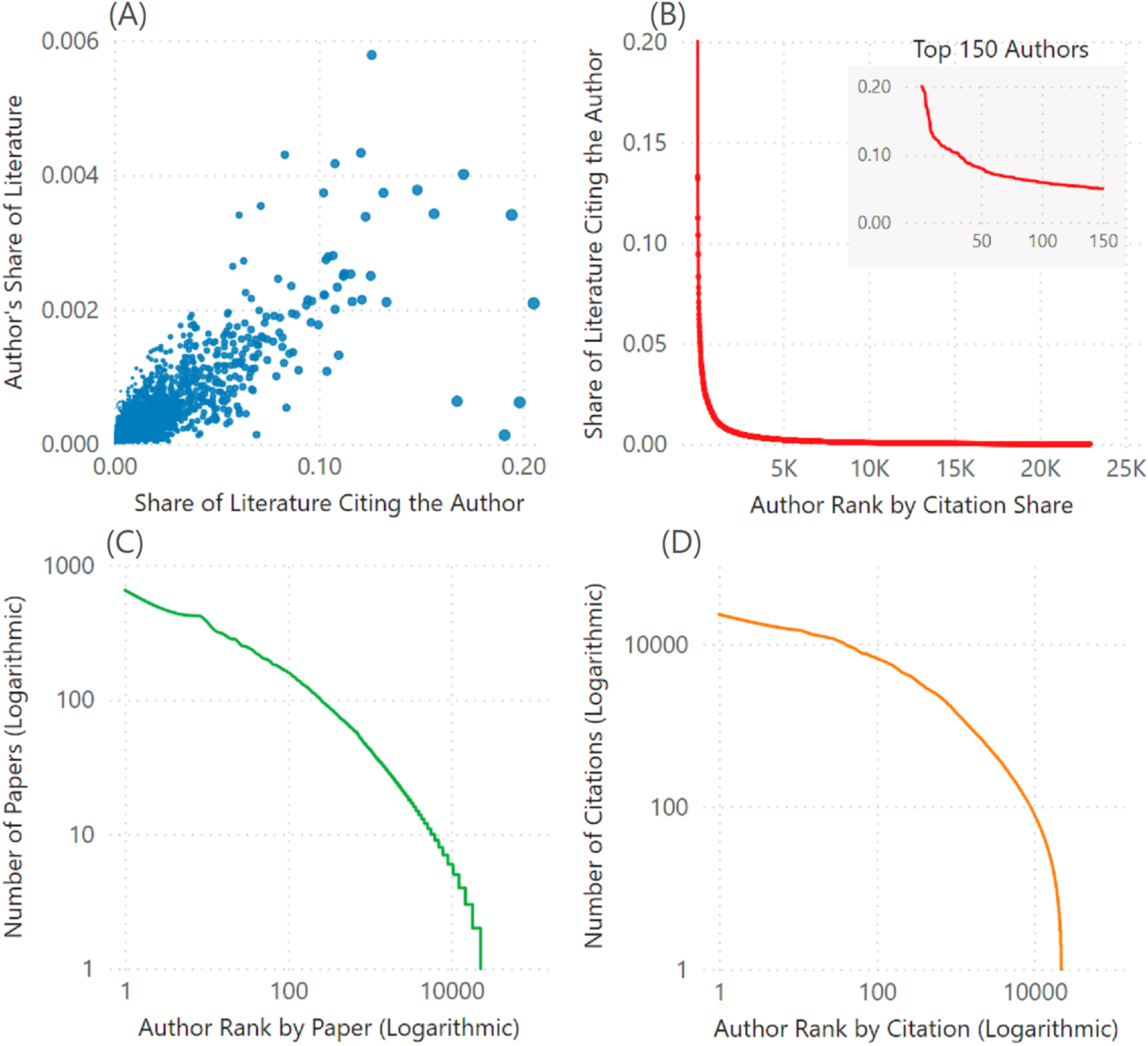
Paper and citation distribution by author. (A) Scatterplot of the fraction of literature citing authors vs. published by authors (p < 2.2e-16, Pearson’s correlation test, n = 22941 authors). (B) The citation reach of authors, all the top 150 (approximately top 1%) (C) Paper distribution by author among those with 2+ papers (D) Citation distribution by author among those with 2+ papers

We next asked to what extent the most productive and highly cited authors in the field could call into question the integrity of later articles that cited and potentially built on that work. The fraction of the AD literature and the number of citations was highly correlated (r = 0.84, p < 2.2e-16, Pearson’s correlation test, n = 22,941 authors, Figure 3A). Of the 22,941 authors who published multiple first-- or last-author AD papers, the average fraction of the literature citing an author was 0.3% (Figure 3B). Less than 1% of these authors were cited by more than 5% of the literature (Figure 3, n = 112,720). In the top 150 authors, the average citation reach of their papers was 7.8% of the literature (n = 112,720). While these highly-cited authors pose a higher risk if they engage in fraudulent activity, on average, 92% of the literature (n = 112,720) in the field does not cite their work. It is important to note that for the literature to be at risk to this extent, researchers engaging in misconduct would need to engage in undetected fraudulent activity in all their papers, and all citations would have to represent causal knowledge transfer rather than ancillary citations (Hoppe et al., 2022). This means that our measures constitute an upper bound on the amount of corruption of the scientific literature that could occur through knowledge transfer documented through citations. Lower frequencies of misconduct or the presence of ancillary citations (Hoppe et al., 2023) would necessarily reduce the meaningful citation reach of such work.

To better understand the potential impact of hypothetical author-level misinformation published in the scientific literature, we visualized the citation reach of randomly selected individual authors cited by approximately 2% of the AD literature (Figure 4). This is a comparable figure to the fraction of the literature citing the author who was called into question in Science (Weiland, 2022). The likelihood of this literature being corrupted by individual authors varies across author groups, where the most influential authors in terms of citation impact undoubtedly have the broadest citation reach. The field appears generally robust against the potential of fraudulent conduct by the vast majority of individual authors. Our results suggest the relatively limited impact of most highly cited authors in the field of AD research and highlight the importance of a diverse and robust knowledge base for advancing scientific understanding. It also raises awareness of the potential risks associated with fraudulent activity but acknowledges that the majority of the literature is not significantly affected by such misconduct.

**Figure 4.**
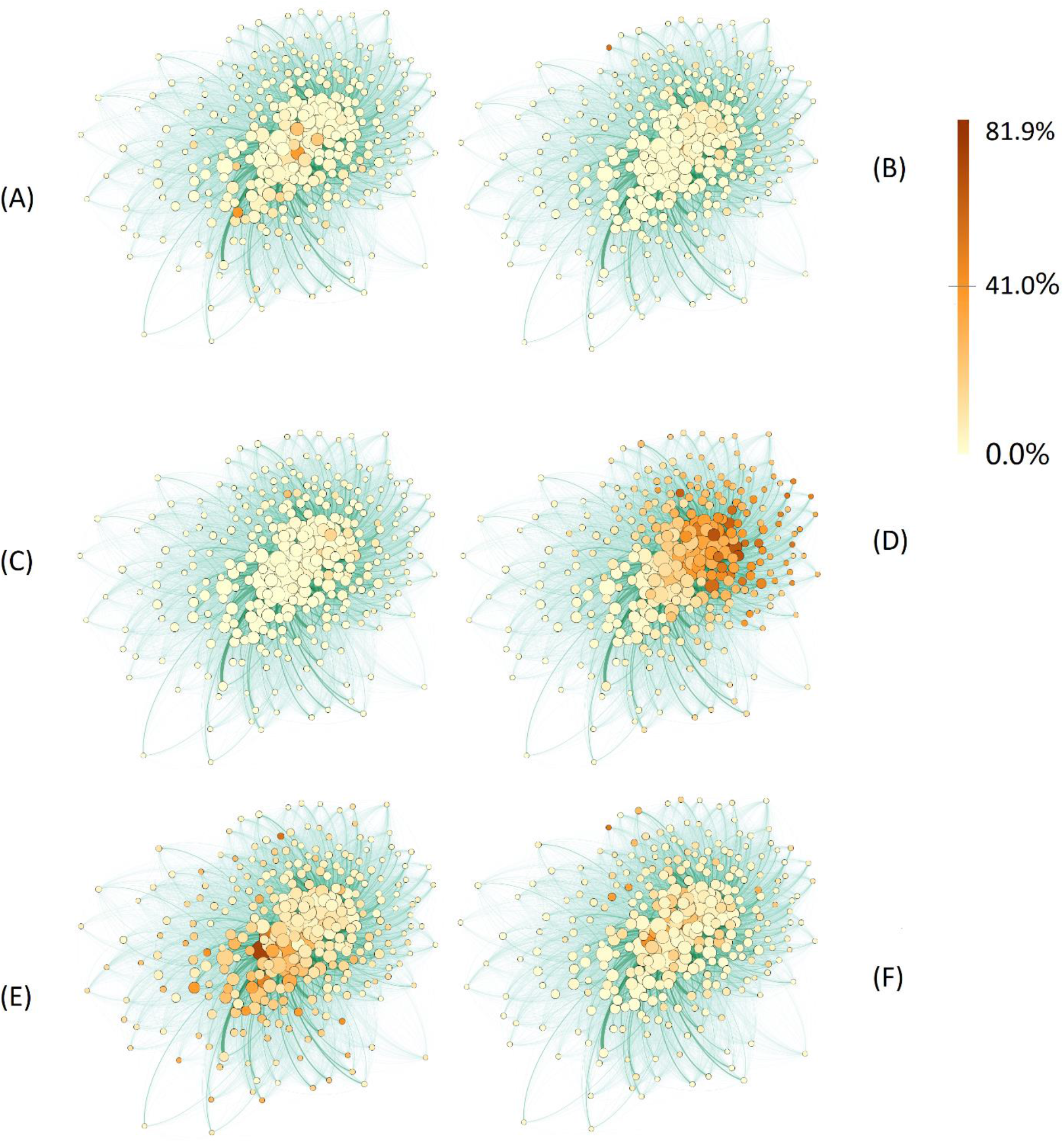
Examples of citation reach by individual authors. Each node represents an AD paper cluster (Methods). The color of each node is proportional to the percent of literature in that node that could in principle be contaminated by an individual scientist if they engaged in research misconduct. (A-C).Citation reach of authors who collectively reach about 2% of the literature, comparable to the example cited in the Science investigation. (D) Citation reach of the most highly cited author in this literature. (E, F) Citation reach of randomly selected authors from the top 250 (∼1%) of citations in this literature.

## Discussion and Conclusions

With questions about research integrity on the rise, there is a growing concern about assessing the risk of malicious actors in specific research fields (Kozlov, 2023; Weiland, 2022). In this study, we examined how fraudulent research practices in publications and among authors could affect the broader field of AD research. Our findings suggest that the risk of individual scientists jeopardizing the integrity and reliability of the majority of the literature in this field is low. Specifically, only a small percentage of pathway citation links, 0.01% (n = 2,051,603), have a direct citation relationship (shortest path=1). Additionally, only a small proportion of publications in the investigated AD research core corpus have been retracted or are currently under further investigation. However, our study highlights the importance of monitoring the integrity of the most productive authors, as they can reach a significant portion of the field, accounting for up to 7.8% of the AD research literature citations (n = 2,051,603) for the top 150 authors and around 21% for the most highly cited author in the field. We also explored the potential impact of indirect citations, which refer to references-of-references in biomedical research articles. The analysis revealed that a considerable number of papers cited both the immediately preceding work and its antecedent, indicating a culture in AD research of acknowledging important preceding world. This suggests that indirect citations are unlikely to transmit problematic information.

The AD study in question has unquestionably been influential. Yet, the amyloid hypothesis, a cornerstone of AD research, traces its origins back to a pivotal study in the 1980s (Glenner & Wong, 1984), which isolated and sequenced the amyloid beta peptide from brain plaques in AD patients. This hypothesis posits that accumulations of amyloid beta protein in brain plaques contribute to neurotoxicity and downstream patient cognitive decline. Subsequent papers (Beyreuther & Masters, 1991; Hardy & Allsop, 1991; Hardy & Higgins, 1992; Selkoe, 1991) further solidified this hypothesis as a plausible mechanism for contributing to AD. This suggests that amyloid deposition may play a central role in the etiology of AD. The hypothesis has greatly influenced the understanding and exploration of AD and has contributed to recent amyloid beta clearing drugs to treat the disease.

Although the amyloid hypothesis has been disputed, proponents have emphasized evidence from human genetic studies supporting the role of the ABeta peptide in AD (Selkoe & Cummings, 2022; Selkoe, 1991; Selkoe & Hardy, 2016). Early evidence from patients with Down’s Syndrome who normally harbor an extra copy of the Amyloid Precursor Protein (APP), suggested that they may suffer from AD symptoms because of its overexpression. Subsequent work showed that Down’s Syndrome patients who do not harbor an extra copy may not experience AD symptoms (Prasher et al., 1998). Duplication of the APP gene outside Down’s Syndrome appears to be associated with autosomal dominant early-onset AD (Rovelet-Lecrux et al., 2006). A mutation (A673T) in the APP gene that decreases the production of a form of ABeta peptides thought to drive AD seems to be protective against AD symptoms (Jonsson et al., 2012). Competing hypotheses posit the involvement of the protein Tau as the primary driver of AD (Brier et al., 2016). Importantly for this study, a comparatively small fraction of the body of AD work seems to draw upon the finding called into question by the Science investigation (Weiland, 2022), which reported a potentially important oligomeric form of ABeta that appears at 56 kDa on a Western blot. Whether one is more persuaded by the evidence about ABeta or Tau, the predictive power of the amyloid hypothesis as a whole does not seem to depend on the presence or absence of a 56 kDa band.

The recent investigation raises broader questions about the vulnerability of the field to researchers who might intentionally publish incorrect information in the literature. Given the accumulated influence the amyloid hypothesis commanded prior to 2006, it is not plausible that the possible invalidation of a subsequently published study could call into question the prior body of work. Notably, a seminal review of the amyloid hypothesis after 25 years (Selkoe & Hardy, 2016) did not cite the paper reported on in *Science*. Drugs based on this hypothesis have had mixed and highly controversial records to date (Kepp et al., 2023). Despite that fact, amyloid plaque-clearing drugs have recently been authorized by the United States Food and Drug Administration (FDA) to treat AD (Reiss et al., 2023). Although the amyloid hypothesis is the most prominent in the Alzheimer’s literature, it drives only 16% of AD drug development (Selkoe & Cummings, 2022; Selkoe & Hardy, 2016).

One critical property of science is that, in the long run, it is self-correcting. This means that scientific knowledge is always subject to revision and refinement based on new evidence, and any errors or inaccuracies can be corrected through further investigation and testing. In fact, the scientific method itself is designed to ensure that hypotheses are continually tested and refined, with new observations and data being used to challenge and modify existing theories. When it comes to unreliable or flawed published research, these self-correcting mechanisms work by either drawing attention to suspicious research outcomes from the citable literature, a process known as retraction, or by systematically ignoring irreproducible research. Retracting flawed research helps limit future citations to the literature (Azoulay et al., 2015). However, retraction cannot completely offset the negative impact of questionable research (Hsiao & Schneider, 2021). Preventing suspicious research from being published is valuable. It is, therefore, crucial that the scientific community continues to uphold high standards of research integrity and transparency to minimize the occurrence of retracted papers (Schneider et al., 2022). Therefore, it is vital for researchers to be vigilant in their citation practices and to carefully evaluate the quality and reliability of the articles in their research.

Our study provides quantitative insights into the potential risks of fraudulent research practices in the field of AD research. While the risk of individual scientists jeopardizing the integrity and reliability of the majority of the literature in this field is generally low, certain aspects, such as highly cited papers and authors, warrant attention. By maintaining high standards of research integrity and transparency, the scientific community can minimize the impact of unreliable or flawed research and ensure the advancement of knowledge in the field of AD research.

Additionally, the framework of analysis employed in this study is suitable for assessing the potential risks of untrustworthy publications or individual malicious actors’ conduct on knowledge production and science advance. Assessing the risk of problematic research and malicious actors in science is complex and challenging. We evaluated the potential risks of questionable research by analyzing the likelihood of individual suspicious papers and researchers contaminating the community through citation relationships. This framework could be conveniently adopted for assessing the risk of suspicious papers and authors in other research fields and domains.

Although this study provides valuable insights, it is essential to recognize that there are limitations to the research that should be taken into account. Our analysis was limited to citation relationships within the Alzheimer’s research community by definition. It is possible that other fields and areas may also draw upon published Alzheimer’s research, which was not included in this study.

Furthermore, the extent to which findings from this single field generalize to other fields and the larger scientific enterprise requires further investigation. Finally, it is an open question how multiple malicious actors publishing simultaneously, either in coordination or independently, could affect information transmission in the scientific community. Given the prevalence of scientific collaboration, future research may investigate this aspect for additional insights.

## Data and Methods

### Data sources

We relied on multiple data sources for this study, including PubMed and Open Citation Collection. To generate a knowledge network of AD research, we began with the National Library of Medicine’s Medical Subject Heading codes (*Medical Subject Headings*, 2022). These are assigned by expert curators to papers indexed in PubMed. We augmented these with the small number of papers mentioning AD in the title or abstract but that were not tagged with this code (< 5% of the total). To analyze the citation network structure of this literature, we used the National Institutes of Health’s Open Citation Collection (Hutchins, Baker, et al., 2019; iCite et al., 2019). Finally, we downloaded author publication profiles from the PubMed Knowledge Graph (Xu et al., 2020).

### Data and processing

To identify the corpus of AD research, we queried PubMed via its API in August 2022. We used the query “Alzheimer Disease[mh] OR Alzheimer[title]” to capture papers that have either been tagged with the AD Medical Subject Heading keyword by curators at the National Library of Medicine or that mention the disease prominently in the article title. Based on these PubMed Identifiers (PMIDs), we cross-referenced this set of articles with citation metadata from the NIH iCite database, including the NIH Open Citation Collection citation graph (Hutchins, 2021; Hutchins, Baker, et al., 2019; Hutchins, Davis, et al., 2019; Hutchins et al., 2016; iCite et al., 2019). To incorporate author-level data, we matched article identifiers to the disambiguated author profiles found in the PubMed Knowledge Graph (Xu et al., 2020). Measurements were taken from distinct samples of citations, paper, or authors, where appropriate for the analysis.

### Author filtering

After generating summary statistics of the entire dataset, we focused our analysis on authors who had published at least one first- or last-author paper at some point in their career. This is because these two authorship positions in biomedical research often signal a prominent role in the given research study. Authors who had only published middle-author papers were excluded, yielding 75,730 who had published at least one first- or last-author AD paper in their career. Most of these authors, however, published only one first- or last-author AD paper in their career, calling into question whether they were focused on AD per se or published a single paper ancillary to their career. Therefore, we focused on the authors who published at least two first- or last-author AD papers, yielding 22,941 AD researchers.

### Clustering and citation path calculations

Visualizing large citation graphs is challenging, so to facilitate visualization of the reach of individual authors, we generated clusters of papers. We used the Leiden clustering algorithm (Traag et al., 2019). Such clustering approaches yield semantically related groups of publications (Klavans et al., 2020; Rahkovsky et al., 2021). The cluster area indicates the number of papers, and shading indicates the fraction of papers within a cluster that cited a given author. For our path length analysis, we used the *igraph* R package (Csárdi & Nepusz).

## Data availability statement

Data needed to evaluate the conclusions of the paper are available at Figshare: https://figshare.com/s/31f7eaf38f888f94d11a

## Code availability statement

Code used to generate the conclusions are available in GitHub: https://github.com/HutchinsLab/AD_risk_analysis

## Notes

### Competing Interest Statement

The authors have declared no competing interest.

